# Integrative Transcriptomic Analysis Identifies Novel Mitochondrial Gene Targets in Parkinson’s Disease

**DOI:** 10.1101/2025.08.20.671402

**Authors:** Anusree Damodaran, Sanskriti Goswami, Sonia Verma

**Affiliations:** Division of Neuroscience and Ageing Biology, CSIR-Central Drug Research Institute, Lucknow, UP, India; Academy of Scientific and Innovative Research (AcSIR), Ghaziabad 201002, India

**Keywords:** Parkinson’s disease, Mitochondria, Substantia Nigra, Dopaminergic Neurons, Neurodegeneration, Novel Therapeutic Targets

## Abstract

Parkinson’s disease (PD) involves the progressive loss of dopaminergic (DA) neurons within the substantia nigra (SN) region of the midbrain, although the precise molecular processes driving this degeneration are still not fully understood. This research investigates the expression patterns of genes associated with mitochondrial function in the SN and DA neurons of individuals with PD, aiming to uncover new potential therapeutic targets. Two independent RNA sequencing datasets, GSE7621 and GSE8397 (GPL-96), retrieved from the GEO database, were analyzed to identify mitochondria-related genes that are differentially expressed in the SN of PD patients. Gene Ontology and pathway enrichment analyses were also performed to gain insight into the molecular mechanisms involved. To validate our findings, we utilized an additional dataset, GSE49036. We also examined the altered expression of these mitochondrial-related genes in DA neurons using RNA-seq data from GSE169755, which includes DA neurons isolated from the SN of both PD patients and healthy controls. Finally, the proposed hypothesis was tested experimentally using an *in vitro* model of PD. This integrative analysis across multiple datasets reveals previously unrecognized mitochondrial gene candidates implicated in PD pathogenesis and highlights their potential as targets for therapeutic intervention.

## Introduction

Parkinson’s disease (PD) is the world’s second most common neurodegenerative disorder, presenting with progressive motor symptoms—such as resting tremor, rigidity, bradykinesia, and gait disturbances—as well as a spectrum of non-motor features including psychiatric, cognitive, and sleep disturbances [1, 2]. These symptoms collectively impair daily functioning and quality of life. The global burden of PD is rising rapidly, with recent projections estimating over 25 million cases by 2050—a 112% increase from 2021—driven primarily by population aging and growth and posing a significant public health challenge, especially in East Asia and middle-income regions [3].

PD is marked by the progressive loss of dopaminergic (DA) neurons in the substantia nigra (SN) and the accumulation of Lewy bodies—intracellular inclusions primarily composed of aggregated α-synuclein—which ultimately leads to reduced dopamine levels in the striatum [4]. Despite advances in symptomatic therapies, such as Levodopa and decarboxylase inhibitors, current treatments are limited by declining efficacy and adverse effects and do not halt disease progression [5–7]. This highlights the critical need to uncover new therapeutic targets that directly intervene in the fundamental mechanisms driving the disease.

Among the diverse pathogenic processes implicated in PD, mitochondrial dysfunction has emerged as a central contributor to neurodegeneration [8]. DA neurons are particularly susceptible to mitochondrial impairment due to their high metabolic demands and limited regenerative capacity [9]. Evidence from neurotoxin models, genetic studies implicating mitochondrial genes, and observations of disrupted mitochondrial dynamics and increased oxidative stress in PD brains all point to mitochondrial dysregulation as a key driver of neuronal loss [10–12]. Moreover, interactions between α-synuclein aggregates and mitochondrial membranes create a pathogenic feedback loop that exacerbates mitochondrial dysfunction and oxidative damage [13, 14].

Given the pivotal role of mitochondria in PD pathophysiology, there is growing interest in targeting mitochondrial pathways for disease modification and neuroprotection [15–17]. Small molecules modulating mitochondrial biogenesis, dynamics, mitophagy, and redox balance are under investigation, but a deeper understanding of mitochondria-related gene alterations in vulnerable neuronal populations is needed to inform precision therapies [18–21].

In this context, our study systematically investigates mitochondrial gene expression alterations in the SN and DA neurons of PD patients using integrative transcriptomic analyses across multiple independent datasets. By identifying and validating novel mitochondria-associated gene candidates, we aim to elucidate molecular mechanisms underlying selective neuronal vulnerability in PD and provide a foundation for developing mitochondria-targeted therapeutic strategies that can halt or reverse disease progression in PD. Using a similar strategy, we recently identified novel ferroptosis-related genes altered in the SN and DA neurons of PD patients [22]. These findings underscore the utility of integrative bioinformatics in uncovering PD-related molecular mechanisms and support the current focus on mitochondrial dysfunction as a complementary pathogenic axis.

## Methods

### Acquisition and Analysis of Data

The datasets were obtained from the publicly accessible NCBI Gene Expression Omnibus (GEO) database (http://www.ncbi.nlm.nih.gov/geo), and the selected datasets (GSE7621, GSE8397 (GPL-96), GSE49036, and GSE169755) were analyzed using GEO2R (http://www.ncbi.nlm.nih.gov/GEO/geo2r/) as mentioned in our previous study [22–26].

In brief, datasets were identified using the keyword ‘Substantia Nigra’ and subsequently filtered to include only human samples and sequencing types classified as either expression profiling by array or high-throughput sequencing. For SN-specific RNA-seq analysis, we further selected datasets with RNA-seq performed on postmortem SN tissue from PD patients and controls, a sample size ≥10, and GEO2R analysis availability. The datasets GSE7621, GSE8397 (GPL-96 and GPL-97), GSE20159, GSE20292, and GSE49036 met these criteria and were included for differential gene expression analysis [23-25, 27-29]. For DA neuron-specific analysis, datasets were chosen if sequencing was performed on RNA from DA neurons isolated from SN, with a sample size ≥10 and GEO2R compatibility. Datasets GSE20141 and GSE182622 were selected [28, 30].

The differentially expressed genes (DEGs) were identified using GEO2R (p ≤ 0.05, |log_2_Fold-change (FC)| ≥ 0.5). To ensure disease relevance, only SN datasets (GSE7621, GSE8397 [GPL-96], GSE49036) showing significant reduction in tyrosine hydroxylase (TH) mRNA—a hallmark of PD—were included for SN analysis [31, 32]. As GSE20141 and GSE182622 did not show reduced TH expression in DA neurons, we also included GSE169755 (sample size <10), demonstrating this characteristic reduction. A summary of the key details for these datasets is provided in **Table 1**.

**Table 1:**
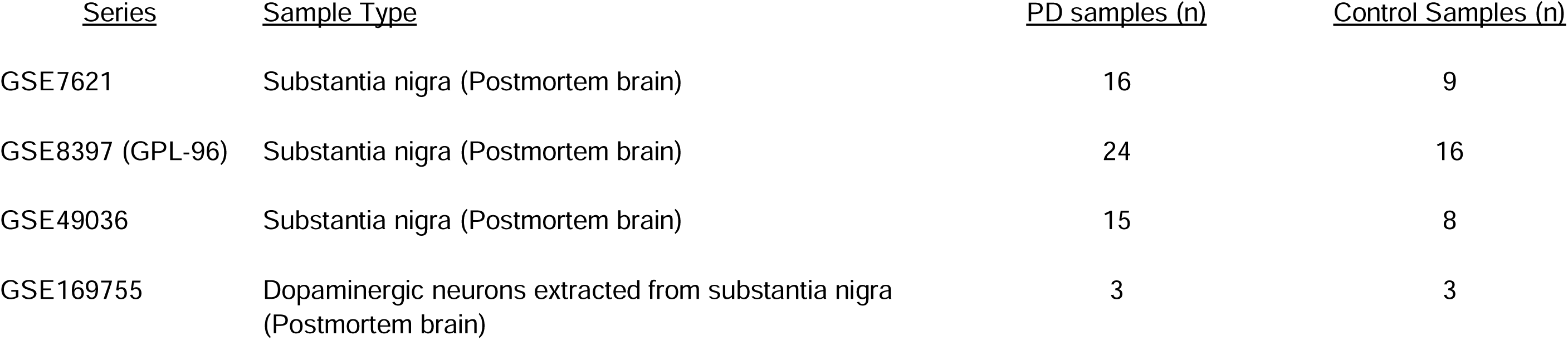
Summary of Transcriptomic Datasets Analyzed in This Study.

| <u>Series</u> | <u>Sample Type</u> | <u>PD samples (n)</u> | <u>Control Samples (n)</u> |
| --- | --- | --- | --- |
| GSE7621 | Substantia nigra (Postmortem brain) | 16 | 9 |
| GSE8397 (GPL-96) | Substantia nigra (Postmortem brain) | 24 | 16 |
| GSE49036 | Substantia nigra (Postmortem brain) | 15 | 8 |
| GSE169755 | Dopaminergic neurons extracted from substantia nigra (Postmortem brain) | 3 | 3 |

### Identification of cDEGs and cDEG_mt_

Common differentially expressed genes (cDEGs) across the datasets were identified and illustrated using Venn diagrams created with the online tool Venny 2.1.0 (http://bioinfogp.cnb.csic.es/tools/venny/index.html).

To identify mitochondria-related genes among the cDEGs (<u>cDEG_mt_</u>), we utilized MitoCarta3.0, a comprehensive database dedicated to a mammalian mitochondrial protein [33]. A comprehensive list of human mitochondria-associated proteins was obtained from the database and is presented in **Supplementary Table 1**. This list was then intersected with the identified cDEGs using Venny 2.1.0, resulting in a subset of cDEG_mt_.

### Functional enrichment analysis of cDEG_mt_

Functional enrichment analysis of Gene Ontology (GO) terms and Reactome pathways associated with cDEG_mt_ was performed using the Database for Annotation, Visualization, and Integrated Discovery (DAVID; https://david.ncifcrf.gov/) [34]. Enrichment was considered significant for GO terms and pathways with p-values ≤ 0.05. The analysis results were visualized as bubble plots using the ‘ggplot2’ package in RStudio (version 1.3.959; https://rstudio.com/) [35].

### Analysis of the Protein-Protein Interaction (PPI) network

A PPI network was generated using the GeneMANIA plugin in Cytoscape (version 3.8.0; http://cyto-scape.org/) [36]. The analysis applied a minimum combined interaction score threshold of ≥ 0.4 to ensure reliable interactions. In the resulting PPI network, nodes represented proteins encoded by the genes of interest, while edges denoted the predicted or known interactions between these proteins.

### Prediction of cDEG_mt_-Transcription Factors Interaction

To predict transcription factors that might interact with cDEG_mt_, we used the NetworkAnalyst platform (https://www.networkanalyst.ca/) and the JASPAR database [37, 38]. Analysis of the resulting cDEG_mt_-transcription factor network revealed key regulatory transcription factors modulating the expression of cDEG_mt_ in PD.

### Prediction of cDEG_mt_-Drug Association

Associations between cDEG_mt_ and potential therapeutics were explored using the NetworkAnalyst platform, which integrates data from the DrugBank database [39]. Identifying drugs linked to cDEG_mt_ may facilitate drug repurposing efforts and expedite the discovery of novel treatment options for PD.

### Maintenance of Cell Culture

SH-SY5Y human neuroblastoma cells (ATCC, Cat. No. CRL-2266) were cultured in DMEM/F12 medium (Gibco, Cat. No. 11320033) supplemented with 10% fetal bovine serum (Gibco, Cat. No. 10270106) and 1% penicillin-streptomycin (Gibco, Cat. No. 15070063). Cultures were maintained at 37°C in a humidified incubator with 5% CO_₂_. The growth medium was refreshed every two days, and cells were subcultured once they reached ∼70% confluency [22, 40].

### Treatment with 1-methyl-4-phenyl 1,2,3,6 tetrahydropyridine (MPTP)

A 10 mM stock of MPTP (Sigma-Aldrich, Cat. No. M0896) was dissolved in dimethyl sulfoxide (DMSO; Sigma-Aldrich, Cat. No. D2650). The stock was diluted in culture medium for cell treatment to achieve a final concentration of 400 μM MPTP, and cells were exposed for 24 hours. Media containing 0.1% DMSO was vehicle control [22, 41].

### Isolation of RNA and Quantitative Reverse Transcription Polymerase Chain Reaction (qRT-PCR)

Total RNA was isolated using the RNeasy Mini Kit (Qiagen, Cat. No. 74104) following the manufacturer’s protocol. Subsequently, one µg of RNA was reverse transcribed into complementary DNA (cDNA) using the cDNA Reverse Transcription Kit (Promega, Cat. No. A5000) following the provided instructions. The reverse transcription reaction was carried out at 42°C for 60 minutes, followed by enzyme inactivation at 70°C for 5 minutes. qRT-PCR was performed using SYBR Premix Ex Taq™ (TaKaRa Bio Inc., Cat. No. RR42OA) according to the manufacturer’s protocol. The amplification protocol included an initial denaturation at 95°C for 30 seconds, followed by 40 cycles of 95°C for 5 seconds and 60°C for 20 seconds. GAPDH was used as the internal control, and relative gene expression was calculated using the 2^−ΔΔCt method. The primer sequences (Origene) utilized in this study are provided in Table 2.

**Table 2:**
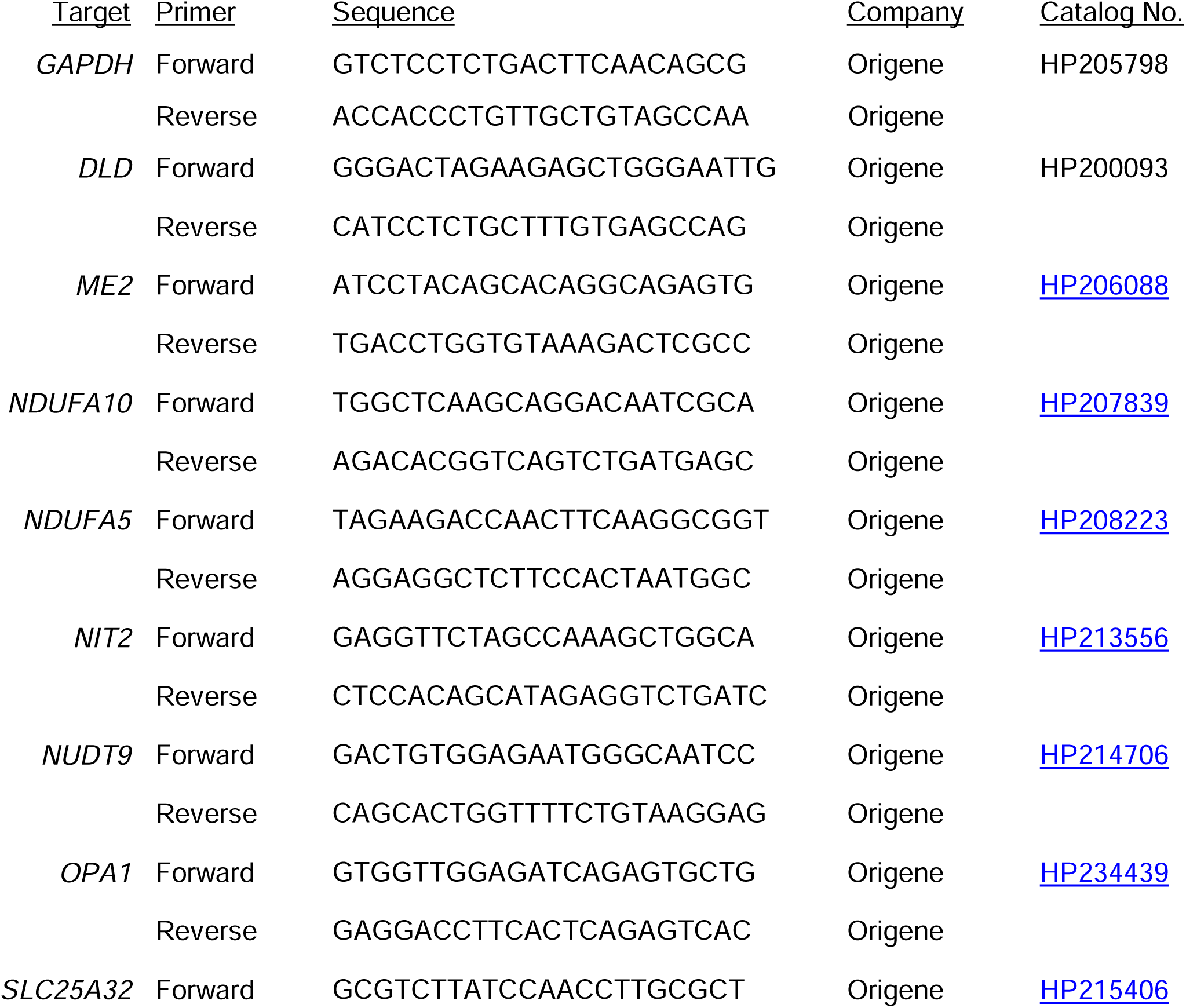
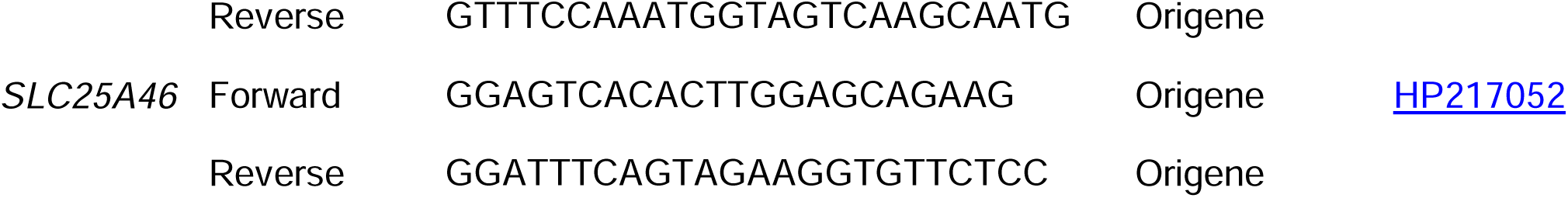
Sequence of Primers used for qRT-PCR.

### Statistical Analysis

Statistical analysis of qRT-PCR results from three independent experiments was performed using GraphPad Prism 8 (GraphPad Software, La Jolla, CA, USA). Data are expressed as mean ± standard deviation (SD). Student’s t-test was used to determine statistical significance, with p-values ≤ 0.05 considered statistically significant.

## Results

### Differential Gene Expression Analysis in SN-PD

A more substantial decrease in *TH* mRNA expression was detected in SN samples from PD patients in datasets GSE7621 and GSE8397 (GPL96) compared to those in GSE49036. This led us to prioritize these two datasets for the initial identification of mitochondrial-associated gene dysregulation in SN-PD while reserving GSE49036 for subsequent validation (**Supplementary Figure 1**).

Using GEO2R with thresholds of p ≤ 0.05 and |log_₂_FC| ≥ 0.5, GSE7621 yielded 3,027 DEGs (1,761 upregulated, 1,266 downregulated in SN-PD vs SN controls) (**Figure 1A and 1C**) and GSE8397 (GPL96) identified 1,490 DEGs (498 upregulated, 992 downregulated in SN-PD compared to SN controls) (**Figure 1B and 1C**).

**Figure 1:**
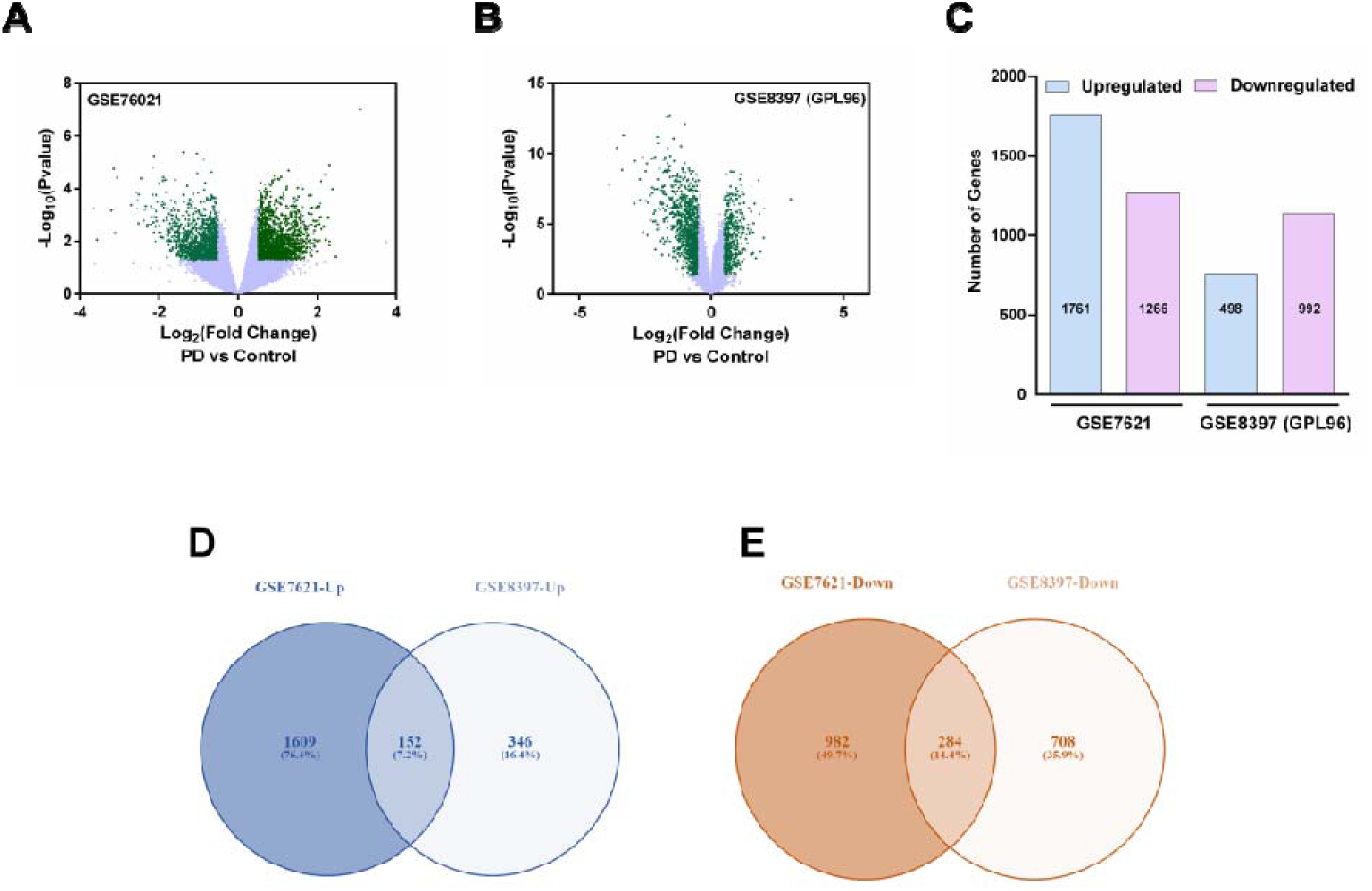
Comparison of gene expression changes in SN-PD versus SN-control using GEO datasets (GSE7621 and GSE8397 (GPL-96)). (A) Volcano plot of dataset GSE7621. (B) Volcano plot of dataset GSE8397 (GPL96). Green dots represent differentially expressed genes with p ≤ 0.05 and |log_2_Fold-change| ≥ 0.5. (C) Bar graph showing the total number of upregulated and downregulated genes (p ≤ 0.05 and |log_2_Fold-change| ≥ 0.5) in SN-PD versus SN-control in each dataset. (D) Venn diagram comparing upregulated genes between datasets GSE7621 and GSE8397 (GPL-96). 152 genes were upregulated in both datasets. (E) Venn diagram comparing downregulated genes between datasets GSE7621 and GSE8397 (GPL-96). 284 genes were downregulated in both datasets. Substantia Nigra-SN; Parkinson’s Disease-PD

Venn analysis revealed 152 consistently upregulated and 284 consistently downregulated genes (**Figure 1D and 1E**). The convergence of 436 cDEGs across independent datasets strongly implicates these genes in PD pathogenesis and provides a focused set of candidates for downstream investigation.

### Identification of mitochondria-associated genes in SN-PD

To explore the association of mitochondrial dysfunction in the SN during PD, we initially retrieved a curated list of 1,136 mitochondria-associated genes from the MitoCarta3.0 database (**Supplementary Table 1**). We then cross-referenced this gene set with the 436 cDEGs. This intersection, visualized using a Venn diagram, revealed 23 cDEG_mt_: five upregulated (*SPR, CYP27A1, SLC25A37, NUDT9*, and *EFHD1*), and 18 downregulated (*OPA1, DLD, MECR, HSDL2, NDUFA5, IDH2, NDUFA10, NIT2, ME2, RMDN1, SLC25A46, YME1L1, SLC25A32, PARL, ACSL6, NIPSNAP1, LYRM9*, and *MTPAP*) in the SN-PD patients across both datasets (**Figure 2A and 2B**). The heatmap (**Figure 2C**) depicts their expression profiles, and detailed gene information is provided in **Table 3**.

**Figure 2:**
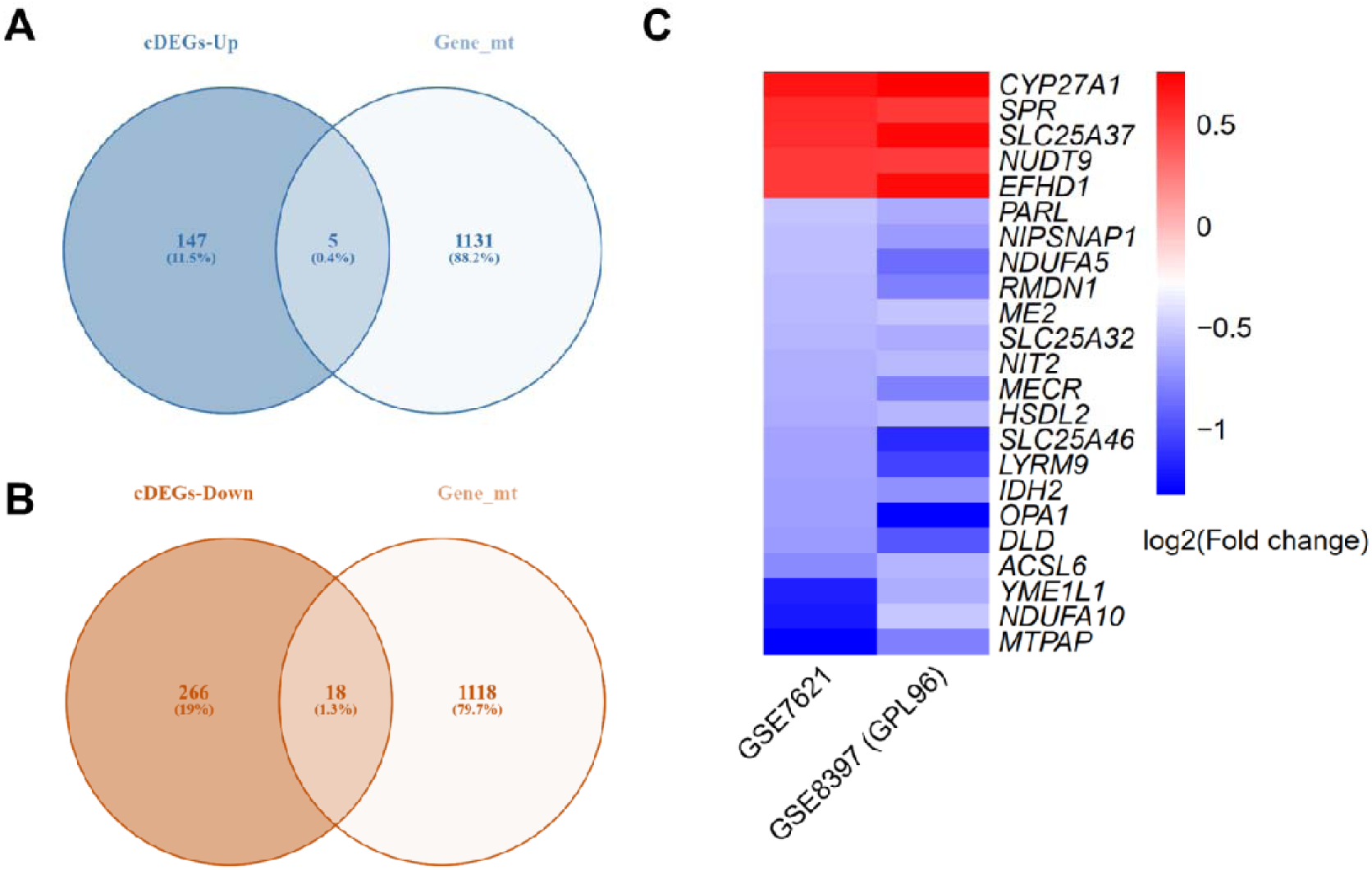
Identification of mitochondria-associated genes among the differentially expressed genes common to GSE7621 and GSE8397 (GPL-96). **A**: Venn diagram showing the overlap between upregulated cDEGs (cDEGs-Up) and the Gene_mt_ acquired from the MitoCarta3.0 database. Five genes were common between cDEGs-Up and Gene_mt_. **B**: Venn diagram showing the overlap between downregulated cDEGs (cDEGs-Down) and the Gene_mt_. 18 genes were common between cDEGs-Down and Gene_mt_. **C**: Heatmap showing the differential expression of the 23 cDEG_mt_ (p ≤ 0.05) between SN-PD and SN-control samples across datasets GSE7621 and GSE8397 (GPL96). Common differentially expressed genes-cDEGs; Mitochondria-associated genes-Gene_mt_; Common differentially expressed mitochondria-associated genes-cDEG_mt_

**Table 3:** Functional Details of cDEG_mt_.

| <u>Gene</u> | <u>Full Name</u> | <u>Function</u> | <u>Reference</u> |
| --- | --- | --- | --- |
| *SPR* | Sepiapterin Reductase | Catalyzes the final step in the biosynthesis of tetrahydrobiopterin, a cofactor essential for producing neurotransmitters like dopamine and serotonin. | [88] |
| *CYP27A1* | Cytochrome P450 Family 27 Subfamily A Member 1 | Involved in converting cholesterol to bile acids, playing a crucial role in cholesterol homeostasis. | [89] |
| *SLC25A37* | Solute Carrier Family 25 Member 37/ Mitoferrin-1 | Functions as a mitochondrial iron transporter and is essential for heme biosynthesis and iron homeostasis. | [90] |
| *NUDT9* | NUDIX Hydrolase 9 | Hydrolyzes ADP-ribose to AMP and ribose-5-phosphate, playing a role in cellular energy metabolism and DNA repair. | [91] |
| *EFHD1* | EF-Hand Domain Family Member D1 | Acts as a calcium-binding protein involved in mitochondrial calcium signaling, influencing processes like apoptosis and energy metabolism. | [92] |
| *OPA1* | OPA1 Mitochondrial Dynamin Like GTPase | Regulates mitochondrial fusion and maintains cristae structure, essential for mitochondrial integrity and function. | [93] |
| *DLD* | Dihydrolipoamide Dehydrogenase | Functions as the E3 subunit in multiple mitochondrial enzyme complexes, playing a vital role in energy metabolism by catalyzing oxidation-reduction reactions. | [94] |
| *MECR* | Mitochondrial Trans-2-Enoyl-CoA Reductase | Catalyzes the final step in mitochondrial fatty acid synthesis, crucial for producing lipoic acid, a cofactor for mitochondrial enzyme complexes. | [83] |
| *HSDL2* | Hydroxysteroid Dehydrogenase Like 2 | Involved in lipid metabolism, particularly in the oxidation of hydroxysteroids, contributing to steroid hormone biosynthesis. | [95] |
| *NDUFA5* | NADH: Ubiquinone Oxidoreductase Subunit A5 | A component of mitochondrial complex I, it plays a role in the electron transport chain, contributing to ATP production. | [96] |
| *IDH2* | Isocitrate Dehydrogenase (NADP(+)) 2, Mitochondrial | Catalyzes the oxidative decarboxylation of isocitrate to α-ketoglutarate in the TCA cycle, producing NADPH and playing a role in cellular metabolism. | [97] |
| *NDUFA10* | NADH: Ubiquinone Oxidoreductase Subunit A10 | Subunit of mitochondrial complex I, involved in electron transfer from NADH to ubiquinone in the respiratory chain. | [96] |
| *NIT2* | Nitrilase Family Member 2 | Functions as an amidase, hydrolyzing specific amide bonds, and may play a role in detoxification processes. | [98] |
| *ME2* | Malic Enzyme 2 | Catalyzes the oxidative decarboxylation of malate to pyruvate, generating NADPH, and is involved in lipid biosynthesis and energy production. | [99] |
| *RMDN1* | Regulator of Microtubule Dynamics 1 | Involved in regulating microtubule dynamics, affecting cell shape, division, and intracellular transport. | [100] |
| *SLC25A46* | Solute Carrier Family 25 Member 46 | Participates in mitochondrial dynamics, particularly in maintaining mitochondrial morphology and distribution. | [101] |
| *YME1L1* | YME1 Like 1 ATPase | A mitochondrial protease involved in the degradation of misfolded proteins and the regulation of mitochondrial morphology. | [102] |
| *SLC25A32* | Solute Carrier Family 25 Member 32 | Transports folate into mitochondria, essential for mitochondrial one-carbon metabolism and nucleotide biosynthesis. | [103] |
| *PARL* | Presenilin Associated Rhomboid Like | A mitochondrial intramembrane protease involved in the regulation of mitochondrial morphology and apoptosis. | [104] |
| *ACSL6* | Acyl-CoA Synthetase Long-Chain Family Member 6 | Activates long-chain fatty acids by converting them to acyl-CoA, playing a role in lipid metabolism and energy production. | [105] |
| *NIPSNAP1* | NIPSNAP Homolog 1 | Involved in mitochondrial quality control and may play a role in the regulation of mitophagy. | [106] |
| *LYRM9* | LYR Motif Containing 9 | Potentially involved in the assembly or stability of mitochondrial protein complexes. | [107] |
| *MTPAP* | Mitochondrial Poly(A) Polymerase | Adds poly(A) tails to mitochondrial mRNAs, playing a role in the stability and translation of mitochondrial transcripts. | [108] |

The consistent identification of these 23 mitochondrial genes across independent datasets highlights the central role of mitochondrial gene dysregulation in PD pathogenesis. These cDEG_mt_ represent promising candidates for further functional studies to elucidate the mechanisms of mitochondrial dysfunction in DA neurodegeneration.

### Functional Enrichment Analysis of cDEG_mt_

We performed GO and pathway enrichment analyses to elucidate the biological significance of the cDEG_mt_. GO biological process analysis revealed significant enrichment in seven key processes: mitochondrial electron transport from NADH to ubiquinone, pyruvate metabolism, mitochondrial cristae formation, mitochondrial fission, 2-oxoglutarate metabolism, negative regulation of cytochrome c release, and mitochondrial fusion (**Figure 3A; Supplementary Table 2**).

**Figure 3:**
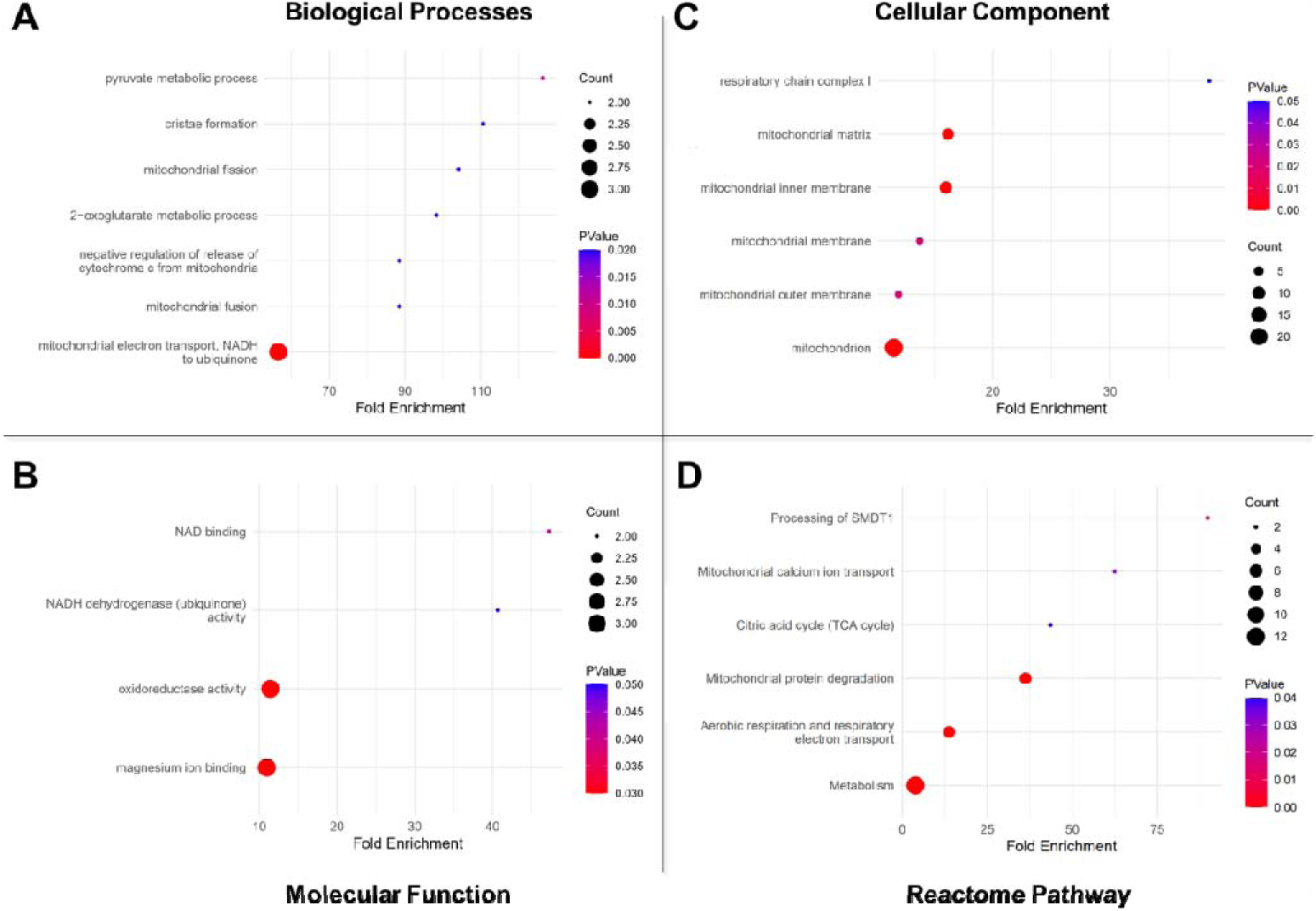
Gene Ontology and Reactome pathway enrichment analysis of the cDEG_mt_ using online software, DAVID. Bubble plots showing the significant GO terms for **(A)** Biological Processes, **(B)** Molecular Function, **(C)** Cellular Component, and **(D)** Reactome Pathway enrichment.

At the molecular function level, four categories were significantly enriched: oxidoreductase activity, magnesium ion binding, NAD binding, and NADH dehydrogenase (ubiquinone) activity (**Figure 3B; Supplementary Table 3**).

GO cellular component analysis further demonstrated that cDEG_mt_ are predominantly localized to the mitochondrion and its subcompartments, including the inner and outer membranes, the matrix, and respiratory chain complex I (**Figure 3C; Supplementary Table 4**).

Pathway enrichment analysis using the Reactome database identified a significant overrepresentation of cDEG_mt_ in pathways related to mitochondrial protein degradation, metabolic processes, aerobic respiration, respiratory electron transport, mitochondrial calcium ion transport, SMDT1 processing, and the tricarboxylic acid cycle (**Figure 3D; Supplementary Table 5**).

In summary, functional enrichment analyses demonstrate that the 23 cDEG_mt_ are intricately involved in multiple aspects of mitochondrial biology, and their dysregulation likely contributes to the multifaceted mitochondrial dysfunction observed in PD.

### PPI Network of cDEG_mt_

To further explore the functional landscape of the cDEG_mt_, we constructed a PPI network using Cytoscape, integrating interaction data from the GeneMANIA database. The resulting network comprised 43 nodes—representing the 23 cDEG_mt_ and 20 additional interacting proteins—and 218 edges, reflecting a high degree of connectivity (**Figure 4A**). This dense interaction pattern suggests that cDEG_mt_ are embedded within shared biological pathways and molecular processes, indicating coordinated roles in mitochondrial regulation relevant to PD.

**Figure 4:**
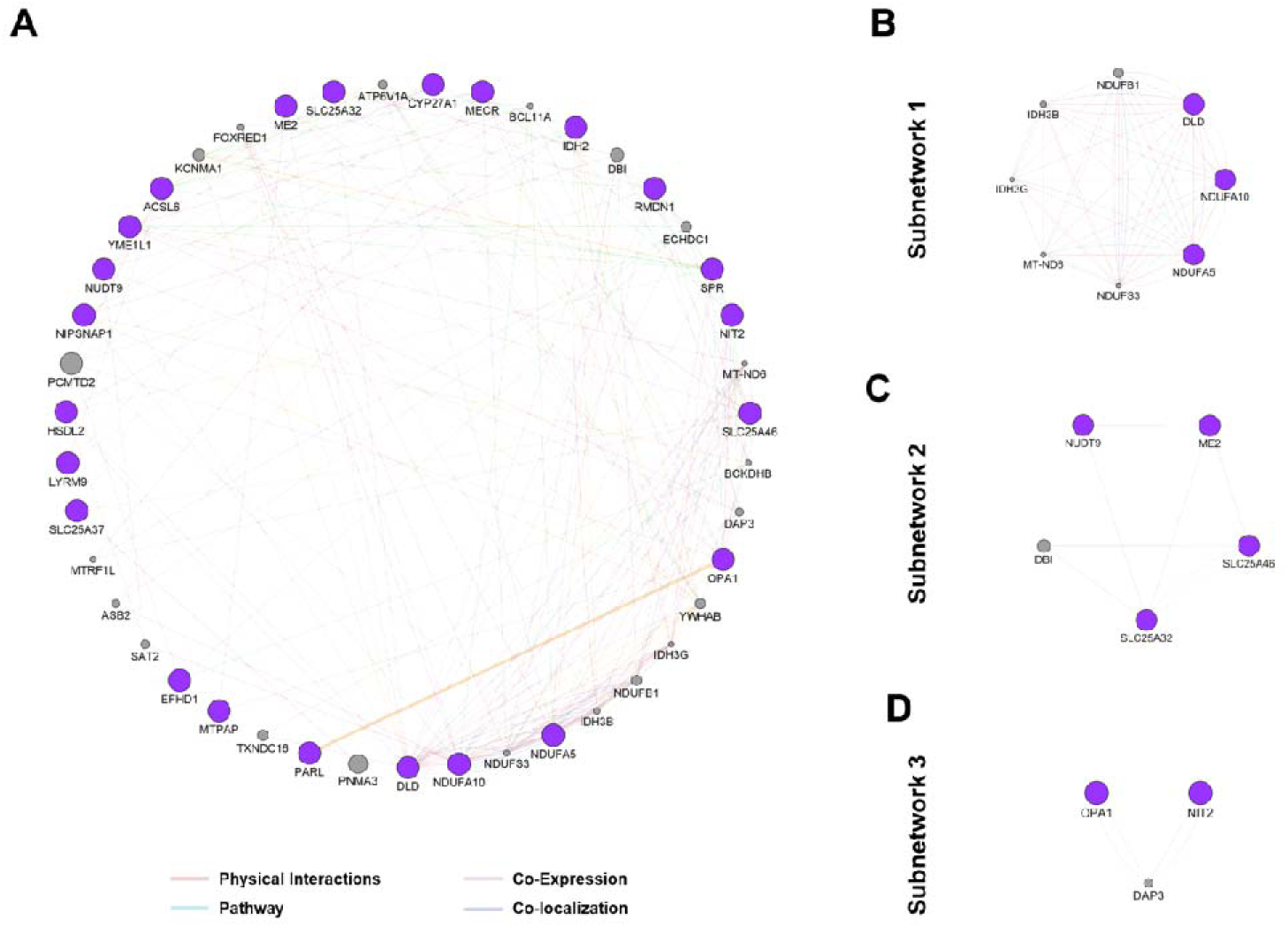
The Protein-Protein Interaction network complex and modular analysis of cDEG_mt_. The purple-colored network nodes are proteins encoded by cDEG_mt_. The edges represent their functional associations. **(A)** Using the GENEMANIA plugin in Cytoscape software, 23 cDEG_mt_ were filtered into the Protein-Protein Interaction network. (**B, C, and D**) The most significant modules were screened using the MCODE plugin in the Cytoscape software.

Network topology analysis identified several cDEG_mt_ as hub genes, characterized by high connectivity, underscoring their potential as central regulators of mitochondrial function. To delineate functional modules within the network, we applied the MCODE algorithm, which revealed three significant subnetworks.

Subnetwork 1: Comprised of eight nodes, with *NDUFA5, DLD*, and *NDUFA10* emerging as hub genes, interconnected by 60 edges (**Figure 4B**).
Subnetwork 2: Included five nodes, *NUDT9*, *ME2*, *SLC25A32*, and *SLC25A46,* identified as hub genes, connected by eight edges (**Figure 4C**).
Subnetwork 3: Contained three nodes, with *OPA1* and *NIT2* as hub genes, linked by five edges (**Figure 4D**).

Identifying these nine hub genes highlights their potential pivotal roles in mitochondrial dysfunction and the broader pathogenesis of PD. Their central positions within the PPI network suggest they may act as key regulatory or structural components within critical mitochondrial pathways. These findings provide a rational basis for prioritizing these hub genes for further experimental validation, to clarify their functional roles in PD, and assess their utility as diagnostic biomarkers or therapeutic targets.

### cDEG_mt_-Transcription Factors Regulatory Networks

Recognizing the pivotal role of transcription factors in orchestrating gene expression, we hypothesized that specific transcription factors may serve as upstream regulators of the identified cDEG_mt_ in PD. We constructed a cDEG_mt_–transcription factor interaction network to investigate this regulatory landscape, comprising 74 nodes (23 cDEG_mt_ and 51 transcription factors) and 138 edges (**Figure 5A**).

**Figure 5:**
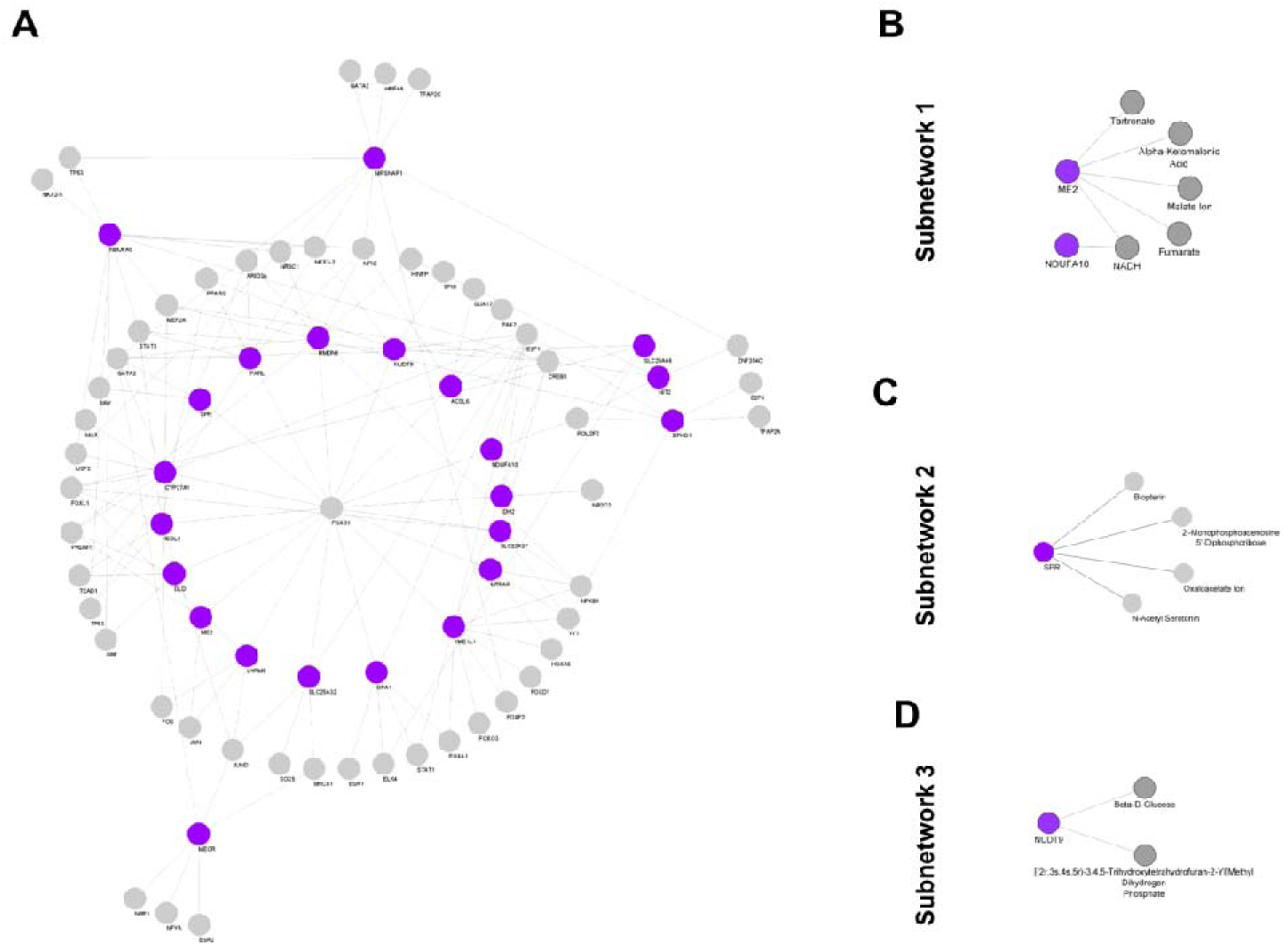
The cDEG_mt_-Transcription Factors and cDEG_mt_–Drug Interaction network created by NetworkAnalyst. The purple-colored network nodes are proteins encoded by cDEG_mt_. **(A)** The The cDEG_mt_-Transcription Factors Interaction network. Gray-colored nodes are transcription factors. (**B**) The The cDEG_mt_-Drug Interaction network. Gray-colored nodes are drugs.

Network analysis revealed that Forkhead box C1 (FOXC1) functions as a central regulatory hub, predicted to modulate the expression of 17 out of the 23 cDEG_mt_. This prominent connectivity suggests that FOXC1 may exert broad regulatory control over mitochondrial gene expression in the SN of PD patients.

This network underscores the potential regulatory influence of specific transcription factors on mitochondrial gene expression in the context of PD and provides new insights into transcriptional mechanisms that may contribute to PD pathogenesis.

### cDEG_mt_-Drug Associations

We conducted a drug interaction network analysis focusing on the cDEG_mt_ to explore potential therapeutic avenues. This analysis revealed three distinct cDEG_mt_–drug subnetworks:

Subnetwork 1: Linked *ME2* to five drugs and *NDUFA10* to one drug (**Figure 5B**).
Subnetwork 2: Associated *SPR* with four drugs (**Figure 5C**).
Subnetwork 3: Connected *NUDT9* with two drugs (**Figure 5D**).

These identified interactions suggest that several cDEG_mt_ are targetable by existing pharmacological compounds, highlighting possible opportunities for drug repurposing in PD.

### Validation of cDEG_mt_ in SN during PD

To independently validate the relevance of the identified cDEG_mt_ in PD, we analyzed the GSE49036 dataset. Differential expression analysis identified 1,897 DEGs (p ≤ 0.05 and |log_₂_FC| ≥ 0.5), comprising 760 upregulated and 1,137 downregulated genes in the SN-PD patients (**Figure 6A and 6B**). Comparing these DEGs with the cDEG_mt_ revealed that nine cDEG_mt_ displayed consistent expression trends across GSE49036, GSE7621, and GSE8397 (GPL96). Notably, *SLC25A37* was consistently upregulated, while *OPA1, ACSL6, NIPSNAP1*, and MT*P*AP were downregulated in the SN-PD patients across all three datasets (**Figure 6C and 6D**).

**Figure 6:**
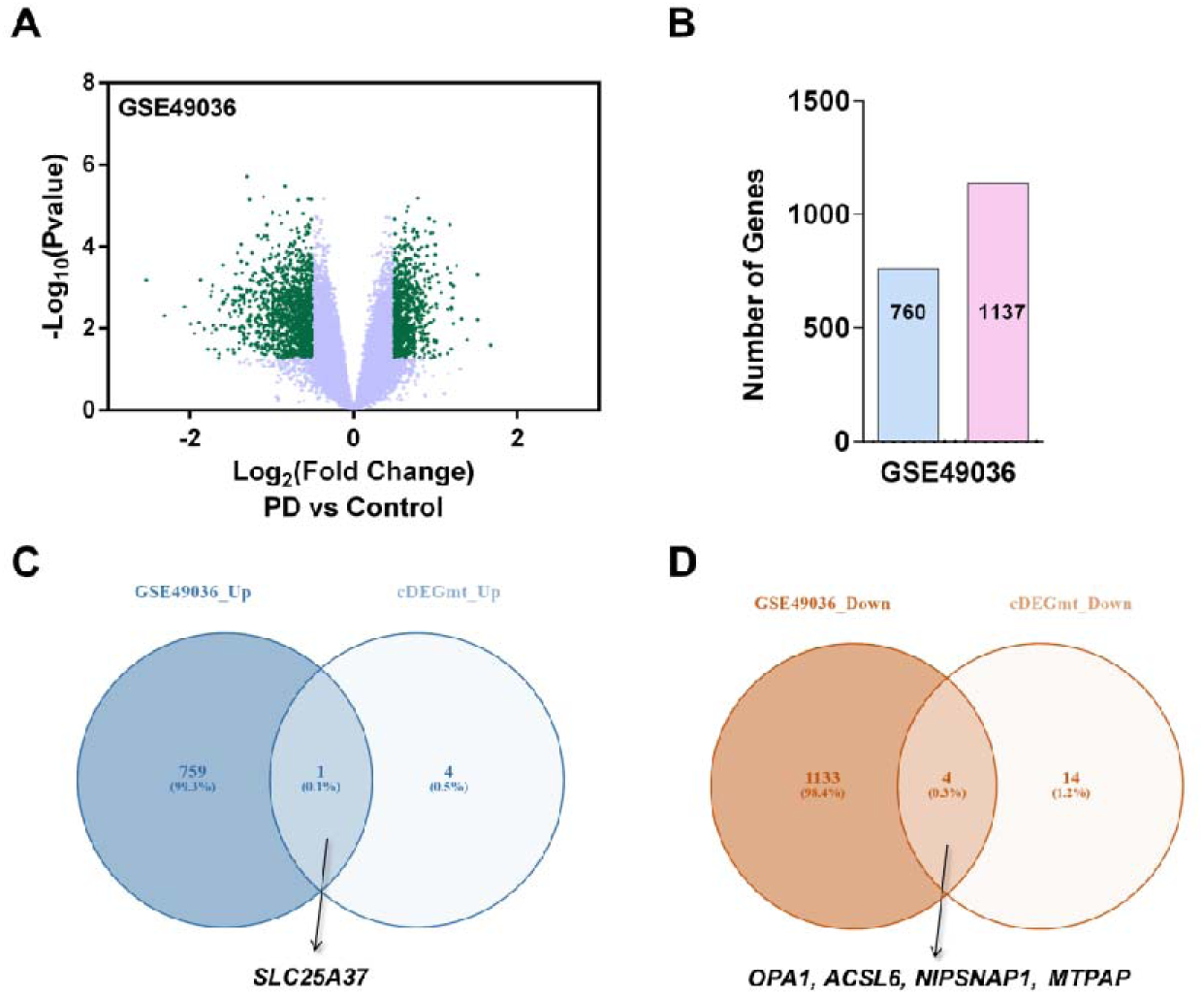
Validation of the cDEG_mt_ in SN-PD using the GEO dataset, GSE49036. (**A)** Volcano plot of dataset GSE49036. **(B)** Bar graph showing the total number of upregulated and downregulated genes (p ≤ 0.05 and |log2Fold-change| ≥ 0.5) in SN-PD versus SN-control in the dataset. (**C)** Venn diagram comparing genes upregulated in SN-PD of GSE49036 with upregulated cDEG_mt_. **D)** Venn diagram comparing genes down-regulated in SN-PD of GSE49036 with downregulated cDEG_mt_.

This concordance in gene expression patterns across independent datasets strengthens the evidence for the involvement of these cDEG_mt_ in PD pathogenesis, particularly concerning mitochondrial dysfunction.

### Validation of cDEG_mt_ in DA neurons during PD

Given the hallmark selective degeneration of DA neurons in PD, we assessed the expression of cDEG_mt_ within PD-related DA neurons using the GSE169755 dataset. Differential expression analysis identified 5,158 DEGs (p ≤ 0.05 and |log_₂_FC| ≥ 0.5), with 2,639 genes upregulated and 2,519 downregulated in DA neurons from PD patients compared to controls (**Figure 7A and 7B**).

**Figure 7:**
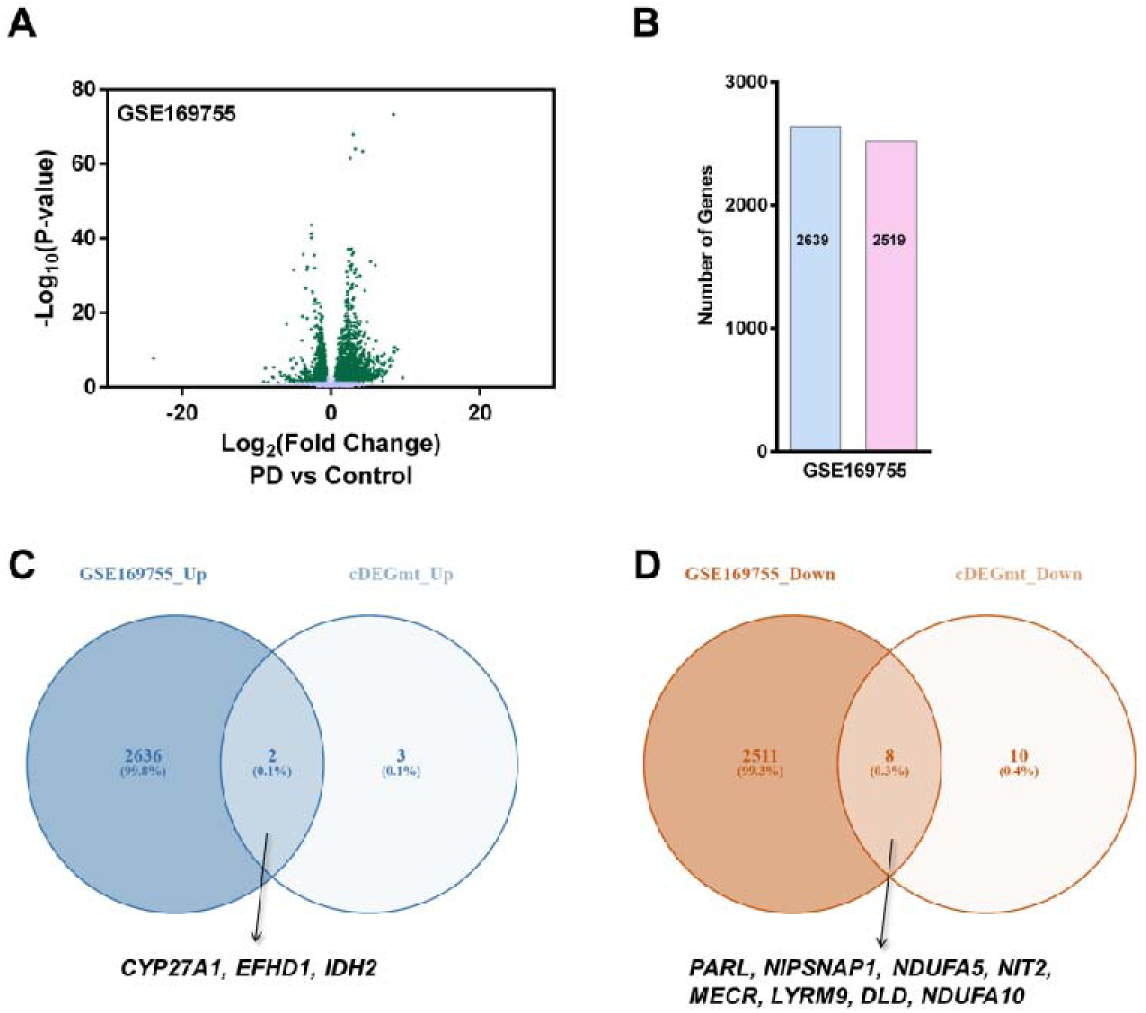
Validation of The cDEG_mt_ in DA-PD using the GEO dataset, GSE169755. (**A)** Volcano plot of dataset GSE169755. **(B)** Bar graph showing the total number of upregulated and downregulated genes (p ≤ 0.05 and |log2Fold-change| ≥ 0.5) in SN-PD versus SN-control in the dataset. (**C)** Venn diagram comparing genes upregulated in DA-PD of GSE169755 with upregulated cDEG_mt_. **D)** Venn diagram comparing genes downregulated in DA-PD of GSE169755 with downregulated cDEG_mt_.

Among the cDEG_mt_, *CYP27A1, EFHD1*, and *IDH2* were consistently upregulated, while *PARL, NIPSNAP1, NDUFA5, NIT2, MECR, LYRM9, DLD*, and *NDUFA10* were downregulated in DA neurons from PD patients (**Figure 7C and 7D**). Importantly, these expression patterns closely mirrored those observed in the SN tissue of PD patients, reinforcing the cellular relevance of these mitochondrial genes.

These findings provide further evidence that dysregulation of cDEG_mt_ is not only a feature of bulk SN tissue but is also present at the level of vulnerable DA neurons. This strengthens the case for these genes as key contributors to mitochondrial dysfunction and neurodegeneration in PD.

### Validating the expression of cDEG_mt_ in the experimental model of PD

To further validate the relevance of cDEG_mt_, we assessed their expression in an established in vitro PD model using SH-SY5Y cells treated with 400 μM MPTP. The qRT-PCR analysis was performed for the nine hub genes. Among these, two genes exhibited significant differential expression in the MPTP-treated cells compared to vehicle-treated controls. Specifically, *NDUFA10* was significantly upregulated, while *ME2* was downregulated in the MPTP-treated cells compared to vehicle-treated controls (**Figure 8A and 8B**).

**Figure 8:**
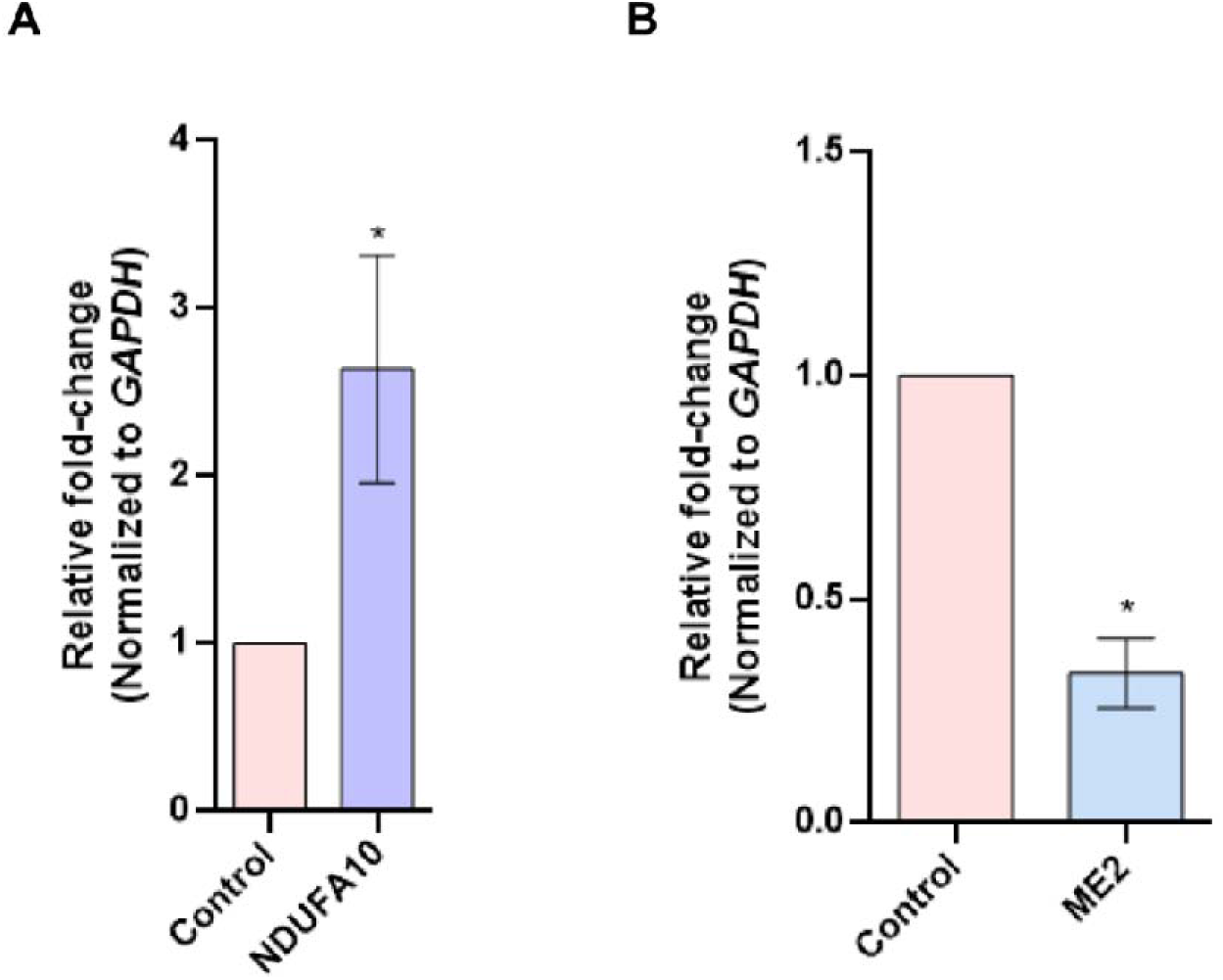
Validating the expression of cDEG_mt_ in the 1-methyl-4-phenyl 1,2,3,6 tetrahydropyridine-treated SH-SY5Y model of PD using qRT-PCR. (A) Bar graph displaying the over-expression of *NDUFA10* in SH-SY5Y cells treated with MPTP (400 μM) for 24 hours. (B) Bar graph displaying the reduced expression levels of *ME2* in MPTP-treated SH-SY5Y cells. Cells treated with 0.1% DMSO were used as controls. GAPDH was used for normalization. The data represent the mean ±S.D (n=3); * p<0.05.

These results demonstrate that a subset of cDEG_mt_ identified as dysregulated in SN through in silico analyses is also altered in an experimental PD model.

## Discussion

This study employed a multi-omics approach to systematically identify mitochondrial-associated genes dysregulated in the SN and DA neurons of PD patients. Analysis of two independent SN transcriptomic datasets (GSE7621, GSE8397) identified 436 cDEGs, of which 23 were mapped to the MitoCarta3.0 mitochondrial database. Multi-cohort validation using SN (GSE49036) and single-nucleus DA neuron (GSE169755) datasets revealed conserved dysregulation patterns for key cDEG_mt_. Experimental validation in an MPTP-induced SH-SY5Y neuronal model further confirmed perturbations in many cDEG_mt_, bridging computational predictions with cellular pathophysiology (**Graphical Abstract**). Critically, several cDEG_mt_ reported here are newly linked to PD, expanding the known repertoire of mitochondrial genes implicated in the disease. These results solidify mitochondrial dysfunction as a central driver of PD pathogenesis and unveil novel molecular targets with therapeutic potential, offering actionable insights for developing disease-modifying strategies.

Functional enrichment analyses revealed that the identified cDEG_mt_ are significantly involved in essential mitochondrial processes that are fundamental for maintaining mitochondrial integrity, neuronal energy balance, and the regulation of apoptosis—all of which are frequently disrupted in PD. Molecular function enrichment further highlighted key enzymatic activities such as oxidoreductase and NADH dehydrogenase activity, both central to redox regulation and electron transport chain function. Disruption of these activities is a well-established feature of PD pathogenesis, contributing to impaired mitochondrial bioenergetics and increased oxidative stress [42, 43].

Cellular component analysis demonstrated that these genes are predominantly localized to critical mitochondrial structures, including the inner and outer membranes, the matrix, and respiratory complex I. This spatial distribution aligns with previous reports of mitochondrial structural and functional abnormalities in PD brains, further supporting the relevance of these cDEG_mt_ to disease pathology [42, 43].

Additionally, Reactome pathway enrichment analysis underscored the involvement of cDEG_mt_ in mitochondrial protein degradation, respiratory electron transport, calcium homeostasis, and the tricarboxylic acid (TCA) cycle. These pathways are vital for maintaining mitochondrial homeostasis and neuronal viability, and their disruption has been closely linked to neurodegeneration in PD [42, 43].

The PPI network analysis, followed by MCODE clustering, identified nine hub genes with high connectivity, underscoring their central roles in maintaining mitochondrial integrity and function in PD. Notably, two of these—*DLD* and *NDUFA5*—have well-established mechanistic links to PD pathogenesis. Dihydrolipoamide dehydrogenase (DLD) is a key component of the mitochondrial α-ketoglutarate dehydrogenase complex. Reduced DLD activity in PD has been shown to elevate oxidative stress and promote neurodegeneration, highlighting its importance in neuronal survival and redox balance [44–46]. *NDUFA5* encodes a core structural subunit of mitochondrial complex I. Downregulation of *NDUFA5* is frequently observed in PD and other neurodegenerative disorders, leading to impaired mitochondrial energy production and increased neuronal vulnerability. This is further supported by studies demonstrating PD-like motor deficits in neuron-specific *Ndufa5* knockout mice and corroborating transcriptomic data from PD models [47–49].

The hub cDEG_mt_ genes *NDUFA10*, *NUDT9*, *ME2*, *SLC25A32*, and *SLC25A46* have not previously been linked to PD, and their identification in this study highlights novel potential contributors to PD pathogenesis. *NUDT9* encodes a mitochondrial ADP-ribose pyrophosphatase that regulates nucleotide homeostasis and redox balance; its dysregulation may compromise respiratory chain efficiency and energy metabolism, thereby exacerbating mitochondrial dysfunction and oxidative stress—central mechanisms in PD neurodegeneration [50, 51]. *ME2* encodes a mitochondrial malic enzyme critical for generating pyruvate and NADH/NADPH, supporting both ATP production and antioxidant defenses; reduced ME2 expression could impair neuronal energy metabolism and increase oxidative vulnerability, potentially accelerating DA neuron loss in PD [52]. SLC25A32 functions as the mitochondrial folate transporter, essential for importing tetrahydrofolate required for one-carbon metabolism, mtDNA synthesis and repair, amino acid and nucleotide metabolism, and NADPH production for redox homeostasis. Disruption of SLC25A32 may impair these mitochondrial folate-dependent pathways, weaken antioxidant defenses, and compromise mitochondrial genome integrity, collectively promoting neurodegenerative processes in PD [53]. SLC25A46 is a mitochondrial outer membrane protein involved in mitochondrial dynamics (fusion and fission) and lipid transfer; while pathogenic mutations in SLC25A46 are known to cause rare neurodegenerative syndromes with parkinsonism, current evidence does not implicate SLC25A46 as a significant factor in typical PD, though its dysfunction could secondarily contribute to parkinsonian features in broader neurological contexts [54, 55]. NDUFA10, an accessory subunit of complex I, is essential for the proper assembly and stability of the holoenzyme. Its phosphorylation by PINK1 directly links it to PD-related mitochondrial dysfunction, and NDUFA10 mutations are implicated in early-onset neurodegenerative disorders such as Leigh syndrome, further highlighting its importance in neuronal survival [56–59]. Together, these findings expand the landscape of mitochondria-associated genes potentially involved in PD, warranting further functional investigation.

Analysis of the cDEG_mt_–transcription factor interaction network identified FOXC1 as the principal regulator of the largest subset of cDEG_mt_, with evidence showing altered FOXC1 protein levels in the SN of PD patients [60]. While the precise role of FOXC1 in DA neuron degeneration remains unclear, FOXC1 is known to regulate genes involved in neurodevelopment, blood-brain barrier maintenance, oxidative stress response, and immune function [61–63]. By influencing these key neuroprotective pathways, FOXC1 may impact PD progression, highlighting its potential as a therapeutic target or biomarker that merits further investigation.

In silico validation using the independent dataset, GSE49036 confirmed the altered expression of several cDEG_mt_ identified across all three SN datasets (GSE7621, GSE8397, and GSE49036). Some of these genes, such as *SLC25A37*, have previously been implicated in PD. SLC25A37 (Mitoferrin-1) encodes a mitochondrial inner membrane protein critical for iron import and heme synthesis; its upregulation in PD models results in excessive mitochondrial iron accumulation, which drives oxidative stress, mitochondrial dysfunction, and neuronal cell death—central mechanisms underlying PD pathogenesis [64].

Notably, the in silico validation also identified cDEG_mt_ genes—*MTPAP, ACSL6, NIPSNAP1*, and *OPA1*—that have not previously been associated with PD, underscoring their novel potential roles in disease progression. *MTPAP* encodes the mitochondrial poly(A) polymerase, essential for mitochondrial mRNA stability and gene expression; although its dysfunction is primarily linked to spastic ataxia type 4, mitochondrial impairment from MTPAP disruption may also contribute to neurodegeneration [65]. NIPSNAP1 is a mitochondrial protein that senses damage and promotes mitophagy by recruiting autophagy machinery; its loss impairs mitochondrial clearance, elevates oxidative stress, and leads to neurodegeneration and motor deficits resembling PD, suggesting it could be a promising therapeutic target [66–68]. ACSL6 is crucial for incorporating neuroprotective DHA into neuronal membranes; its deficiency alters lipid metabolism, increases neuroinflammation, and disrupts dopamine and glutamate homeostasis, all closely related to PD pathology [69–71]. OPA1 maintains mitochondrial cristae integrity and respiratory chain function; its downregulation or mutation impairs ATP synthesis and increases oxidative stress, contributing to syndromic parkinsonism and neurodegeneration and making OPA1 a strong candidate for therapeutic intervention aimed at restoring mitochondrial health in PD [72–77]. Collectively, these findings expand the landscape of mitochondria-associated genes implicated in PD and highlight new avenues for mechanistic studies and therapeutic development.

Analysis of the GSE169755 dataset revealed that several cDEG_mt_ are differentially expressed in DA-PD, including established PD-associated genes such as *CYP27A1* and *PARL*. *CYP27A1* encodes sterol 27-hydroxylase, which generates 27-hydroxycholesterol—a metabolite elevated in PD that promotes α-synuclein aggregation, mitochondrial dysfunction, and neurotoxicity. Genetic deletion of *CYP27A1* in mice reduces α-synuclein pathology and motor deficits, highlighting the therapeutic potential of targeting the CYP27A1–27-OHC axis [78, 79]. *PARL* encodes a mitochondrial protease essential for PINK1 processing and PINK1/Parkin-mediated mitophagy; its downregulation impairs mitochondrial quality control and contributes to PD-related dysfunction, positioning *PARL* as a key therapeutic target [80].

Notably, we report for the first time that *EFHD1, IDH2, NIT2, MECR*, and *LYRM9* are altered in both SN-PD and DA-PD. EFHD1 is a mitochondrial calcium-binding protein involved in mitochondrial dynamics and calcium homeostasis; its upregulation may increase neuronal vulnerability in PD [81]. NIT2 (omega-amidase) detoxifies neurotoxic metabolites, supporting redox balance; reduced NIT2 may elevate oxidative and nitrative stress, accelerating neurodegeneration [82]. MECR is essential for mitochondrial fatty acid synthesis and lipoic acid production; its deficiency impairs respiratory chain function, promotes iron accumulation, and increases susceptibility to ferroptosis, linking MECR dysfunction to PD pathogenesis [83–85]. LYRM9 supports the assembly of mitochondrial respiratory chain complexes and energy production; although direct evidence in PD is limited, its dysfunction could contribute to mitochondrial deficits and neurodegeneration [86]. IDH2 generates NADPH for glutathione regeneration and antioxidant defense; its upregulation in PD likely reflects a compensatory response to oxidative stress, but persistent DA neuron loss suggests this defense is insufficient, emphasizing the need to target mitochondrial redox balance in PD therapy [87]. Together, these findings expand the repertoire of mitochondrial genes implicated in PD and identify new candidates for mechanistic studies and therapeutic intervention.

In summary, this study provides an initial investigation into the expression of mitochondria-related genes in PD, laying the groundwork for future research to elucidate their roles in disease progression and evaluate the potential of mitochondria-targeted therapeutic strategies. Additionally, future studies should focus on delineating the cell-type-specific contributions of cDEG_mt_ in PD. The current identification of mitochondria-related genes builds on our earlier work on ferroptosis-associated gene networks in PD, which were also altered in the SN and DA neurons. Together, these studies illustrate the power of this integrative analytical framework to reveal converging pathways of neuronal vulnerability, offering insight into multiple therapeutic entry points for PD [22].

While this study offers a thorough multi-dataset analysis, certain limitations must be considered. One key limitation is the selection of datasets based on markedly decreased *TH* mRNA expression, which, although relevant to core PD pathology, may introduce bias by emphasizing TH-related mechanisms and potentially overlooking other critical pathways, such as those involving LRRK2. Future studies should consider including datasets reflecting a broader range of PD-related mechanisms. Second, using transcriptomic data from diverse platforms enhances robustness but may introduce method-specific variability, potentially omitting some candidates or causing inconsistencies. Integrating multi-omics approaches and standardizing data acquisition will be important for validation. Third, although validation was conducted using an *in vitro* model, this approach may not fully reflect the intricate pathophysiology of PD *in vivo*. Future studies should incorporate *in vivo* models to provide more physiologically relevant insights. Fourth, while several cDEG_mt_ were identified, the precise molecular mechanisms by which these genes contribute to PD remain unclear. Finally, realizing the therapeutic potential of these target genes will require comprehensive preclinical and clinical investigations, including the evaluation of small-molecule modulators in conjunction with current PD treatments.

## Data availability

The datasets analyzed in this study are publicly accessible through the GEO database [https://www.ncbi.nlm.nih.gov/geo/], with specific accession numbers detailed within the manuscript.

## CRediT authorship contribution statement

A- Investigation; Data curation; Formal analysis; Validation; Visualization; Writing - review & editing.

SG- Investigation; Data curation; Formal analysis; Validation; Visualization; Writing - review & editing.

SV – Conceptualization; Investigation; Data curation; Formal analysis; Methodology; Project administration; Resources; Supervision; Validation; Visualization; Writing - original draft; and Writing - review & editing.

## Abbreviations

cDEGs: Common Differentially Expressed Genes
cDEG_mt_: Common Differentially Expressed Mitochondria-Associated Genes
DA: Dopaminergic
DA-PD: Dopaminergic Neurons in Parkinson’s Disease
DAVID: Database for Annotation, Visualization, and Integrated Discovery
DEGs: Differentially Expressed Genes
DLD: Dihydrolipoamide Dehydrogenase
DMSO: Dimethyl Sulfoxide
DMEM: Dulbecco’s Modified Eagle Medium
FC: Fold-Change
FOXC1: Forkhead Box C1
GAPDH: Glyceraldehyde 3-Phosphate Dehydrogenase
GEO: Gene Expression Omnibus
GO: Gene Ontology
GPL: GEO Platform
MCODE: Molecular Complex Detection
MPTP: 1-methyl-4-phenyl 1,2,3,6 tetrahydropyridine
NCBI: National Center for Biotechnology Information
PD: Parkinson’s Disease
PPI: Protein-Protein Interaction
qRT-PCR: Quantitative Reverse Transcription Polymerase Chain Reaction
SN: Substantia Nigra
SN-PD: Substantia Nigra in Parkinson’s Disease
TH: Tyrosine Hydroxylase

