## Supplementary figures and images for "Integrative Transcriptomic Analysis Identifies Novel Mitochondrial Gene Targets in Parkinson’s Disease"

### Supplementary Figure 1

#### Slide 1
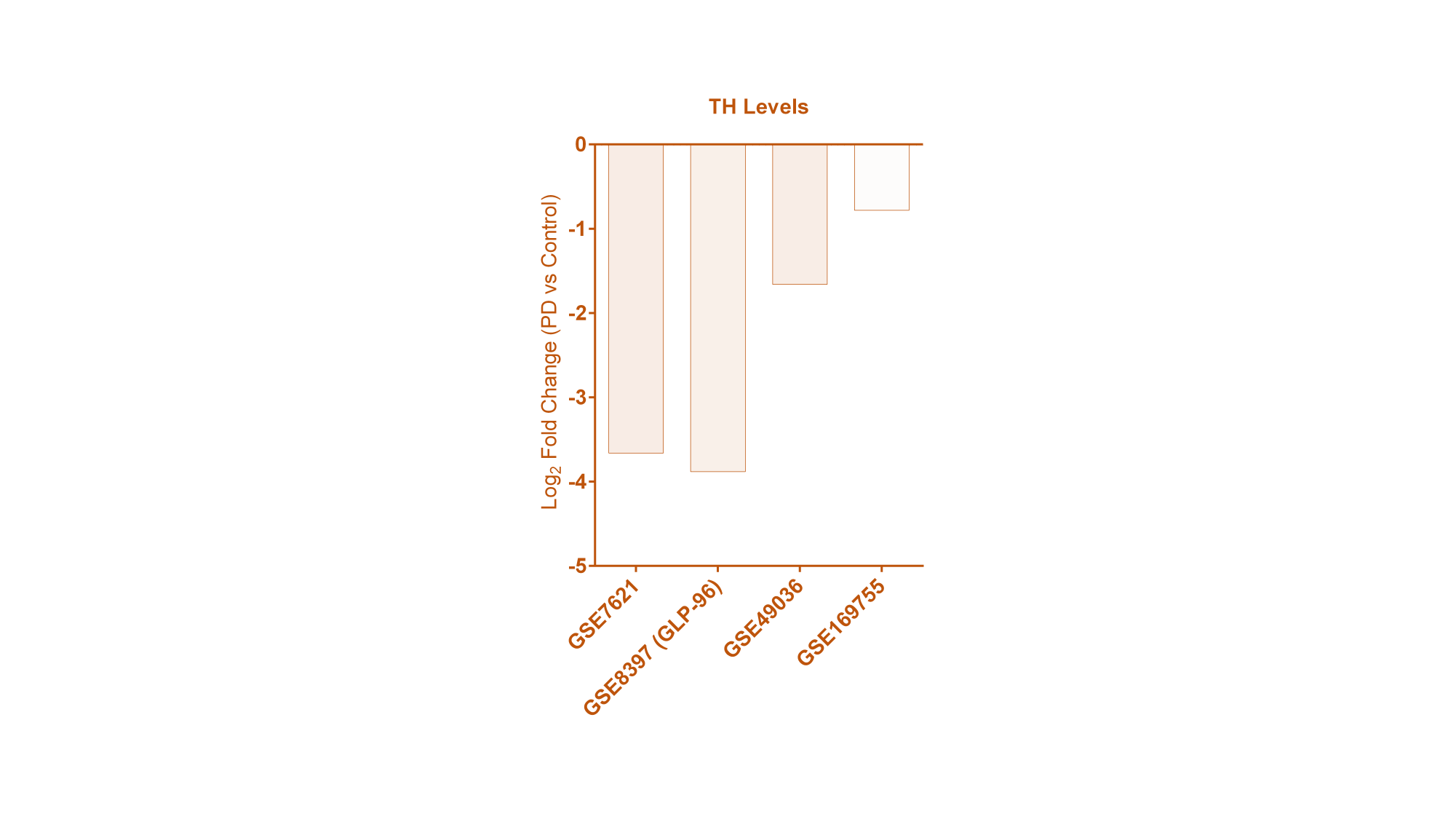
